# C-Jun N-terminal kinase post-translational regulation of pain-related Acid-Sensing Ion Channels 1b and 3

**DOI:** 10.1101/2021.03.18.435937

**Authors:** Clément Verkest, Sylvie Diochot, Eric Lingueglia, Anne Baron

## Abstract

Neuronal proton-gated Acid-Sensing Ion Channels (ASICs) participate in the detection of tissue acidosis, a phenomenon often encountered in painful pathological diseases. Such conditions often involve in parallel the activation of various signaling pathways such as the Mitogen Activated Protein Kinases (MAPKs) that ultimately leads to phenotype modifications of sensory neurons. Here, we identify one member of the MAPKs, c-Jun N-terminal Kinase (JNK), as a new post-translational positive regulator of ASIC channels in rodent sensory neurons. Recombinant H^+^-induced ASIC currents in HEK293 cells are potently inhibited within minutes by the JNK inhibitor SP600125 in a subunit and species dependent manner, targeting both rat and human ASIC1b and ASIC3 subunits but only mouse ASIC1b subunit. The regulation by JNK of recombinant ASIC1b- and ASIC3-containing channels (homomers and heteromers) is lost upon mutation of a putative phosphorylation site within the intracellular N- and the C-terminal domain of the ASIC1b and ASIC3 subunit, respectively. Moreover, short-term JNK activation regulates the activity of native ASIC1b- and ASIC3-containing channels in rodent sensory neurons and is involved in the rapid potentiation of ASIC activity by the proinflammatory cytokine TNFα. Local JNK activation *in vivo* in mice induces a short-term potentiation of the acid-induced cutaneous pain in inflammatory conditions that is partially blocked by the ASIC1-specific inhibitor mambalgin-1. Collectively, our data identify pain-related channels as novel physiological JNK substrates in nociceptive neurons, and propose JNK-dependent phosphorylation as a fast post-translational mechanism of regulation of sensory neuron-expressed ASIC1b- and ASIC3-containing channels that may contribute to peripheral sensitization and pain hypersensitivity.

## Introduction

Chemodetection of noxious and innocuous stimuli by sensory neurons encompasses the detection of a broad range of chemically distinct molecules such as protons, lipids, irritants, toxins or cytokines, often encountered in inflammatory or itch processes (Petho and Reeh, 2012). The various pain-related mediators that are released act on receptors and associated signaling pathways, which are critical in the establishment of pain hypersensitivity (Hucho and Levine, 2007). It includes several protein kinases like protein kinase A or C (PKA, PKC) and also several members of the mitogen-activated protein kinase (MAPK) family like extracellular signal-regulated kinase (ERK), p38 and c-jun N-terminal kinase (JNK) (Obata et al., 2004). JNK is of particular interest regarding its activation in DRG neurons that has been well established in various pain models including migraine (Huang et al., 2016), neuropathic and cancer pain (Simonetti et al., 2014).

Acid-Sensing Ion Channels (ASICs) are critical for the detection of tissue acidosis, which is a hallmark of several painful diseases (Dinkel et al., 2012; Deval and Lingueglia, 2015). At least six subunits are expressed in rodents (ASIC1a, ASIC1b, ASIC2a, ASIC2b, ASIC3 and ASIC4) encoded by four genes, which associate into trimeric functional channels that are widely distributed in the nervous system and all along the pain pathway. Besides protons, ASICs can be modulated by multiple pain-related mediators, including lipids (Smith et al., 2007; Marra et al., 2016), small molecules (Cadiou et al., 2007; Li et al., 2010) and neuropeptides (Askwith et al., 2000). Early work on ASICs in sensory neurons has shown that the ASIC3 subunit is an important player in different pain models (Deval et al., 2008; Yan et al., 2013; Marra et al., 2016). Recent pharmacological and genetic evidences have refined this idea by proposing that ASIC1-containing, and especially sensory neuron-specific ASIC1b-containing channels, can also participate in peripheral pain (Bohlen et al., 2011; Diochot et al., 2012; Diochot et al., 2016; Verkest et al., 2018; Chang et al., 2019).

We combine here functional and pharmacological *in vitro* and *in vivo* approaches to demonstrate that the MAPK JNK is a potent post-translational regulator of the ASIC1b- and ASIC3-containing channels. JNK activation potentiates within minutes the activity of recombinant and native ASIC1b- and ASIC3-containing channels in a species-dependent manner. The effect is dependent of key residues in a putative phosphorylation site within the N-terminal domain of rat, mouse and human ASIC1b, and the C-terminal domain of rat and human, but not mouse, ASIC3. Local activation of JNK *in vivo* in mice evokes a short-term and partially ASIC1-dependent potentiation of the acid-induced cutaneous pain in inflammatory conditions.

## Materials and Methods

### Animals

Experiments were performed on male C57BL/6J mice of 6 to 14 week-old weighting 20-25g and 6-8 week-old male Wistar rats (Charles River Laboratories). Animals were housed in a 12 hours light-dark cycle with food and water available *ad libitum* and were acclimated for at least one week before being subjected to experiments. Animal procedures were approved by the Institutional Local Ethical Committee and authorized the French Ministry of Research according to the European Union regulations and the Directive 2010/63/EU (Agreements E061525 and APAFIS#20260-2019040816489112). Animals were sacrificed at experimental end points by CO_2_ euthanasia.

### Chemicals

For stock solutions, anisomycin (Tocris), SP600125 (Selleckchem) were dissolved in DMSO and TNFα (Peprotech) in saline solution (NaCl 0.9%), and final dilution was made just before the experiments. The final concentration of DMSO never exceeded 0.1% including in control extracellular solution, during patch-clamp experiments, and never exceeded 0.5% including in the vehicle saline solution, during *in vivo* behavior experiments. ASIC-targeting toxins (PcTx1, APETx2 and mambalgin-1) were purchased from Smartox and dissolved in water or saline, with 0.05% Bovine Serum Albumin (BSA) added to the final solution for patch-clamp or behavior experiments in order to avoid non-specific adsorption on vessels and tubings.

### Mutagenesis and site prediction analysis

Point mutants were obtained by recombinant PCR strategies as previously described (Salinas et al., 2009) using the following primers: rASIC1b S59A, forward: 5’ AGGGTGATGACGCACCTAGGGAC 3’, reverse: 5’ GTCCCTAGGTGCGTCATCACCCT 3’. mASIC3 A509T, forward: 5’ CCTCCTACCACTCCCAGTGCT 3’, reverse: 5’ AGCACTGGGAGTGGTAGGAGG 3’. Constructs were subsequently subcloned into pIRES2-EGFP vector using NheI-EcoRI restriction enzymes. All constructs were fully sequenced to confirm the presence of the desired mutation.

Site prediction analysis was carried on intracellular N-terminal and C-terminal sequences of ASIC1a, 1b, 2a, 2b and 3 from different species (mouse, rat and human, Uniprot as source) using Eukaryotic Linear Motif (ELM) resource (Dinkel et al., 2012). Subsequent analysis for putative phosphorylation sites was run using Group Prediction Site software (GPS v5.0) (Xue et al., 2008) with a medium sensitivity threshold in order to refine the list of potential protein kinases.

### Cell culture

HEK293 cells were grown in DMEM medium (Lonza) with 10% fetal bovine serum (Biowest) and 1% peniciline/streptomycine (Lonza) in an incubator at 37°C with 5% CO_2_. One day after being plated at a density of 20,000 cells in 35 mm diameter dishes, cells were transfected with one or several of the following plasmids: pIRES2-rASIC1a-EGFP, pCI-rASIC1a, pIRES2-rASIC1b-EGFP, pCI-rASIC1b, pCI-rASIC2a, pIRES2-rASIC2b-EGFP, pCI-rASIC2b, pIRES2-rASIC3-EGFP, pCI-rASIC3, pCDNA3.1 rTRPV1, pIRES2-EGFP, pIRES2-hASIC1b-EGFP, pIRES2-hASIC3a-EGFP or pIRES2-hASIC1a-EGFP vectors using the JetPEI reagent according to the supplier’s protocol (Polyplus transfection SA, Illkirch, France). 0.5μg of DNA per dish was generally used, except 1μg for homomeric ASIC3. For co-expression of two ASIC subunits, a 1:1 plasmid ratio was used. Fluorescent cells were selected for patch-clamp recordings 2–4 days after transfection. Heteromeric currents were identified by combining several biophysical and pharmacological properties.

DRG neurons were prepared from male C57Bl6J mice (6–14 weeks old) and Wistar male rats (6-8 weeks old) and processed in a similar way, unless mentioned, than previously described (Francois et al., 2013). After isoflurane anesthesia, animals were killed by decapitation and thoracolumbar DRG were collected on cold HBSS and enzymatically digested at 37°C for 40min with collagenase II (Biochrom, 2mg/ml, 235U/mg) for mice and collagenase I (Worthington, 4mg/ml, 370U/mg) for rats and dispase II (Gibco, 5mg/ml, 1,76U/mg) for both. Gentle mechanical dissociation was done with a 1ml syringe and several needles with progressively decreasing diameter tips (18G, 21G and 26G) to obtain a single cell suspension. Neurons were plated on poly-D-lysine/laminin coated dishes with the following medium: Neurobasal-A (Gibco) completed with L-glutamine (Lonza, 2mM final), B27 supplement (Gibco, 1X) and 1% peniciline/streptomycine. One to two hours later, neurons were carefully washed to remove cellular debris and incubated with complete medium. The following additional growth factors were used: Nerve Growth Factor (NGF) 100ng/ml and retinoic acid, 100nM (both from Sigma), Glial Derived Neurotrophic Factor (GDNF) 2ng/ml, Brain Derived Neurotrophic Factor (BDNF) 10ng/ml and Neurotrophin 3 (NT3) 10ng/ml (all three from Peprotech). Patch-clamp recording were done 1-4 days after plating on both “small” (membrane capacitance <40pF) and “large” (>40pF) diameter neurons. Acutely-dissociated mouse DRG neurons were processed the same way as above, except that they were incubated without the additional neurotrophic growth factors listed above and recorded 3 to 10h after plating.

### Patch-clamp recording experiments

Ion currents were recorded using the whole cell patch-clamp technique. Recordings were made at room temperature using Axopatch 700A (Axon Instruments) or 200B (Molecular Devices) amplifiers, with a 2-5 kHz low-pass filter. Data were sampled at 10 kHz using pClamp9 or 10 softwares. The borosilicate patch pipettes (2–7 MΩ) contained the following (in mM): 135 KCl, 5 NaCl, 2 MgCl_2_, 2 CaCl_2_, 5 EGTA and 10 HEPES (pH7.25 with KOH). To investigate regulations by mediators or protein kinases in HEK293 cells and DRG neurons, 2.5mM Na_2_-ATP was used instead of NaCl. The control bath solution contained the following (in mM): 145 NaCl, 5 KCl, 2 MgCl_2_, 2 CaCl_2_, 10 glucose and 10 HEPES (pH7.4 with NaOH). MES was used instead of HEPES to buffer solutions with pH≤6. Cells were voltage-clamped at a holding potential of −60mV (HEK293) or at −80mV for neurons (to minimize potassium current contribution). ASIC currents were activated by a rapid pH drop by shifting one out of eight outlets of a gravity driven perfusion system from a control solution (*i.e.,* pH7.4 or pH8.0) to an acidic test solution. To test the effect of a mediator or a signaling pathway, ASIC currents were repetitively activated every 30s for ASIC1b homomeric current, every 15s for ASIC3 homomeric current, and every minute for ASIC1a homomeric current, these intervals allowing their full reactivation, and drugs were perfused for at least 5 minutes and up to 10 minutes in the pH7.4 control bath solution until their maximal effect was observed. Activation curves were determined by activating whole-cell ASIC currents at a given pH after a 30s perfusion of control pH8.0 solution. Inactivation curves were obtained by maximally activating whole-cell ASIC currents by pH5.0, pH4.5 or pH4.0 after a 30s perfusion at a given pH. The curves of pH-dependent activation and inactivation were fitted by a Hill function: *I* = *a* + (*I*_max_ – *a*)/(1 + (pH_0.5_/[H]*n*)), where *I*_max_ is the maximal current, *a* is the residual current, pH_0.5_ is the pH at which half-maximal activation/inhibition of the transient peak ASIC current was achieved, and *n* is the Hill coefficient. Time constant of current decay were fitted with a single exponential function.

Outside-out configuration was achieved after whole-cell configuration, by slowly pulling out the patch pipette. Intracellular and extracellular medium were the same as above. Cells were then perfused with SP600125 for 5min in the pH7.4 bath solution and a drop to pH5.0 was used to open ASIC channels.

For DRG neurons, action potentials (APs) were evoked in current-clamp mode by 2ms depolarizing current injections of increasing amplitudes (Δ50 or 100pA), and neurons that display a resting membrane potential superior to −40mV and/or no action potential were discarded. Time derivative of membrane voltage was calculated to determine if the falling phase had one or two components (a feature of nociceptive neurons, (Petruska et al., 2000). In voltage-clamp mode, the pharmacological characterization of the ASIC-like peak current was done by applying for 30s the ASIC inhibitory toxins APETx2 (3μM), PcTx1 (20 nM), and mambalgin-1 (1 μM), before the pH drop from resting pH7.4 to pH5.0. Toxin experiments were usually performed with activation of the current from resting pH7.4 to pH5.0. An activating pH of 6.6 (from resting pH7.4) was additionally tested when necessary to improve the functional characterization of the ASIC current. Current was considered as sensitive to a toxin if at least 20% of inhibition was observed. Investigation of the effect of a signaling pathway was then done as described previously on HEK293 cells. 154 mouse DRG neurons from 10 cultures, 49 rat DRG neurons from 4 cultures and 28 acutely-dissociated mouse DRG neurons from 4 cultures were used in the analysis.

### *In vivo* behavior experiments

Acid-induced cutaneous pain was evoked in mice by the intraplantar (*i.pl.*), subcutaneous injection (20 μl, 30-G needle) in the left hindpaw of a pH5.0-buffered solution (NaCl 0.9%, MES 20 mM, pH5.0 with NaOH). To test their local pharmacological effect, drugs were pre-injected 10 minutes before (10μl, 30-G needle) in a non-buffered saline solution (NaCl 0.9%) to allow diffusion within the tissue, then co-injected with the pH5.0 solution. Anisomycin and SP600125 were tested at 500 μM (a higher concentration than the one used in patch-clamp experiments to account for *in vivo*/*in vitro* differences), and mambalgin-1 was tested at 34 μM as previously described (Diochot et al., 2012; Diochot et al., 2016). The final concentrations of DMSO and BSA were 0.5% (anisomycin, SP600125, anisomycin + SP600125, vehicle) and 0.05% (mambalgin-1 containing solutions and related vehicle), respectively.

Mice were placed in mirror boxes where they acclimated for at least 15 min before undergoing a single pH5.0 *i.pl.* injection or the double *i.pl.* protocol (pre-injection followed by co-injection) for pharmacological experiments. Duration of spontaneous pain-related behavior (paw licking, shaking and lifting) was measured over 5 minute-intervals during 15 minutes after the pH5.0 *i.pl.* injection, or from the pre-injection done 10 minutes before in the double *i.pl.* protocol.

Paw inflammation was induced by injection of a 2% carrageenan solution (NaCl 0.9%) in the left hindpaw (20 μl, 25G) 3 hours before measuring the acid-induced cutaneous pain. Mice that did not show a visible red oedema (6 out of 174) and mice that showed more than 15 seconds left hindpaw movements during the 10 min period before the pH5.0 injection (20 out of 169) were excluded in order to avoid non-painful behavior (*e.g.,* grooming) or spontaneous pain from other causes (*e.g.*, wounds).

### Data statistical analysis

Patch-clamp experiments - Data are presented as mean ± SEM along with the individual data points (number of recorded cells in the figure legends). Data from the same cell before and after treatment were analyzed by a paired t-test or a Wilcoxon non-parametric paired test, and two different groups were compared by an unpaired t-test or a Mann Witney non-parametric unpaired test. For multiple comparisons, a one-way ANOVA with Tukey’s post-hoc test for multiple comparisons was performed. *p*<0.05 was considered statistically significant.

Behavioural experiments - Data are presented as mean ± SEM as a function of time, with number of mice written in the figure legends. Data from the same mice were analyzed by a two-way Anova followed by a Dunnett post-hoc paired comparison with data obtained during the 5 min interval preceding the second *i.pl.* injection. Data from different mice were compared by a two-way Anova followed by a Dunnett post-hoc unpaired comparison with the data obtained when the *i.pl.* pH5.0 second injection was realized in the presence of anisomycin. *p*<0.05 was considered statistically significant. Analysis in both patch-clamp and behavioural experiments was performed using GraphPad Prism v4.0 and v6.0 (GraphPad Software, Inc.).

## Results

### Recombinant ASIC1b- and ASIC3-containing channels are positively regulated by short-term JNK activation

Activation of the MAPK JNK has been shown to play important roles in pain, and we investigated the effect of short-term JNK activation on ASIC currents expressed in HEK293 cells transiently transfected with recombinant ASIC subunits. At the holding potential of −60 mV, a rapid drop of the extracellular pH from 7.4 to 5.0 evoked rat ASIC1b (rASIC1b) transient inward current (Fig. 1A). When the cells were incubated several minutes with the MAPK activator anisomycin (10 μM) a small increase in rASIC1b peak current amplitude was sometimes observed but remaining not significantly different from the control current (Fig. 1B). Subsequent incubation with the JNK inhibitor SP600125 (10 μM) for several minutes induced a strong decrease of the rASIC1b current amplitude (Fig. 1A) below the control amplitude, which was consistent with inhibition of a high basal level of JNK activity explaining the lack of significant mean stimulatory effect of anisomycin. A potential non-specific inhibitory effect of SP600125 through direct interaction with the channel was ruled out as the drug had no effect on the rASIC1b current amplitude in outside-out patches (n=3; not shown). To estimate the JNK total effect on the current, we calculated the ratio of rASIC1b peak current amplitude between the maximally JNK-stimulated (*i.e.,* after anisomycin) and the maximally JNK-inhibited (*i.e.,* after SP600125) currents. JNK increased the rASIC1b current amplitude by approximately 50% for every test pH used (+63.2±11.6% at pH6.3, +48.4±7.9% at pH6.0 and +48.0±10.9% at pH5.0) (Fig. 1B). JNK regulation did not change the biophysical properties of rASIC1b, with similar pH_0.5_ values for pH-dependent activation (6.08±0.70 after anisomycin versus 6.05±0.09 after SP600125) as well as pH-dependent inactivation (6.78±0.04 after anisomycin versus 6.80±0.04 after SP600125) (Fig. 1G) and also similar inactivation time constant (Supplementary Fig. 1A).

**Figure 1:**
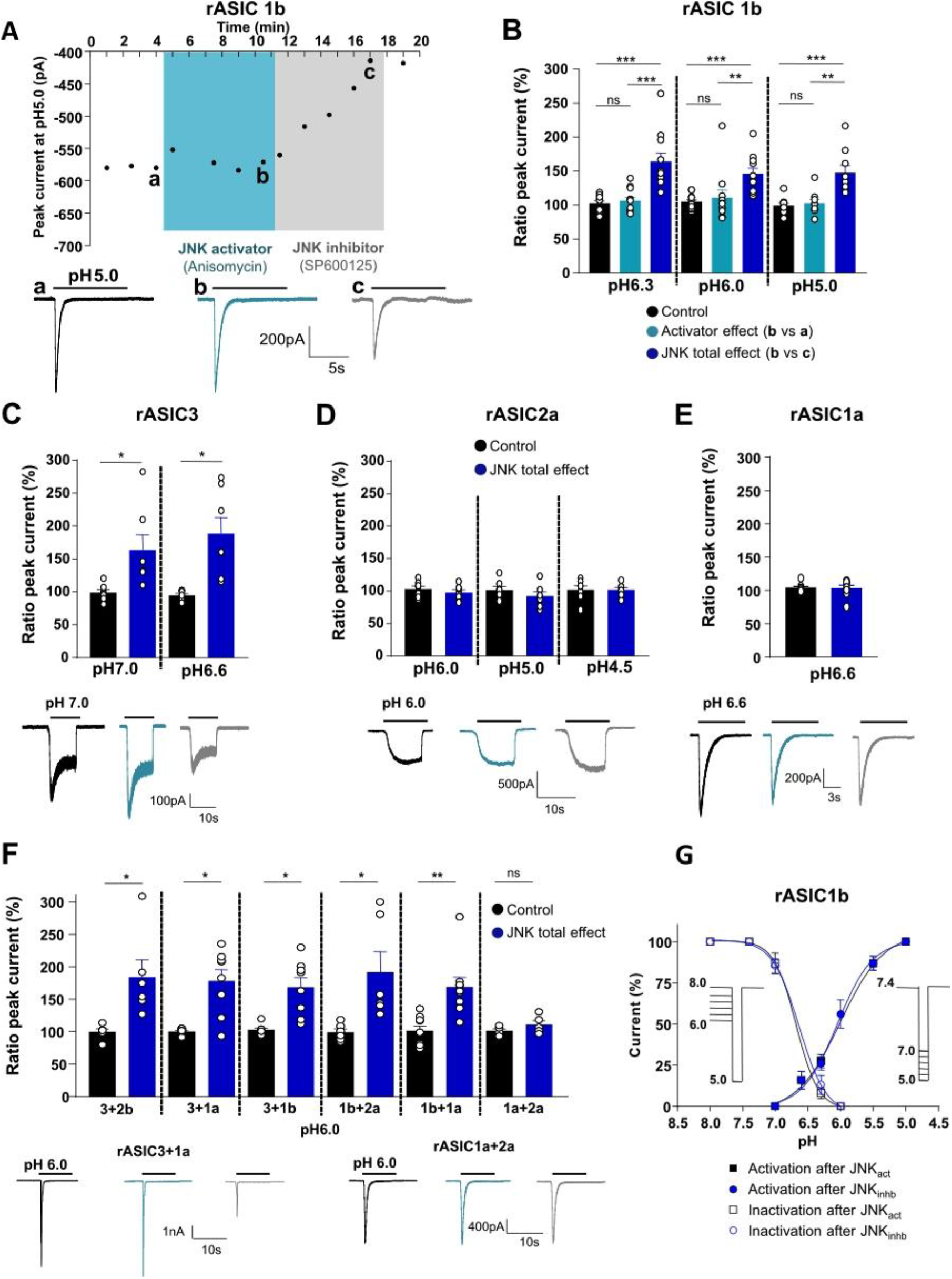
Regulation of recombinant rodent ASIC peak currents by JNK. **A,** Top: Representative time course of whole-cell rASIC1b current peak amplitude recorded at −60mV in HEK293 cells. Currents were activated by a pH drop from 7.4 to 5.0. Anisomycin (50μM) and the JNK inhibitor SP600125 (50μM) were successively applied in the extracellular solution. Bottom: Representative current traces of rASIC1b current in control condition (a, black), after anisomycin application (b, cyan) and after SP600125 application (c, grey). **B,** Potentiation by JNK of rASIC1b current. Ratio (%) of amplitude of H^+^-activated rASIC1b peak current activated at different pHs (conditioning pH7.4), in the three pharmacological conditions shown in A. Individual data points and mean ± SEM are shown (n= 9-11 cells). Control (black) values were calculated from the ratio of the last two control currents, activator effects (cyan) were calculated from the ratio of b/a current amplitudes, and the total JNK effects (blue) were calculated from the ratio of b/c, *i.e.,* the ratio between the current amplitude after anisomycin and the maximally inhibited current amplitude after SP600125 perfusion. One-way ANOVA (F(2, 20)=26.50, p<0.0001) followed by a Tukey post-hoc test, *** p<0.005, **p<0.001. **C,D,E**,**F**, Potentiation by JNK of rASIC3 (C), rASIC2 (D), rASIC1a (E) and heteromeric (F; subunit combinations <u>given below the bar graphs</u>) currents. Ratio (%) of amplitude of H^+^-activated ASIC peak current activated at different pHs (conditioning pH7.4). Mean potentiation by JNK (total effects) was calculated as in B. n=5-9 cells for each channel subtype. Wilcoxon paired test versus control, *, p<0.05, ***, p<0.005. Bottom in C-E: Representative current traces recorded at −60mV in control condition (black), and after anisomycin and SP600125 application (cyan and grey, respectively). **G,** pH–dependent activation and inactivation curves of rat ASIC1b current after anisomycin (black) and SP600125 (blue) application on the same cells (n=4-14). Protocols used for activation and inactivation are shown in inset.

We next explored the effect of JNK in the same conditions on the other ASIC channel isoforms. The amplitude of the peak current flowing through rat ASIC3 (rASIC3) homomeric channels was also significantly increased (+63.4±23.1% at pH7.0 and +88.0±25.0% at pH6.6) (Fig. 1C). On the other hand, currents recorded from rat ASIC1a and ASIC2a (rASIC1a and rASIC2a) homomeric channels were not potentiated for every test pH used (Fig 1D, E). We next tested the effect of JNK on several ASIC heteromers. JNK potentiated the peak current of all the rat ASIC1b- or ASIC3-containing heteromeric channels tested, but not of the rat ASIC1a+2a channels (Fig 1F). The sustained currents associated to ASIC3 (Salinas et al., 2009), *i.e.*, the window current activated by pH close to 7.4, as well as the pH5.0-evoked sustained current, were also potentiated by JNK (+53.4±13.5% for rASIC3 at pH7.0 and +27.4±6.1% for rASIC3+2b at pH5.0) (Supplementary Fig. 1B). The currents flowing through human ASIC1b and ASIC3 channel isoforms were potentiated by JNK in a similar way than their rat counterparts (Fig. 2A,B).

**Figure 2:**
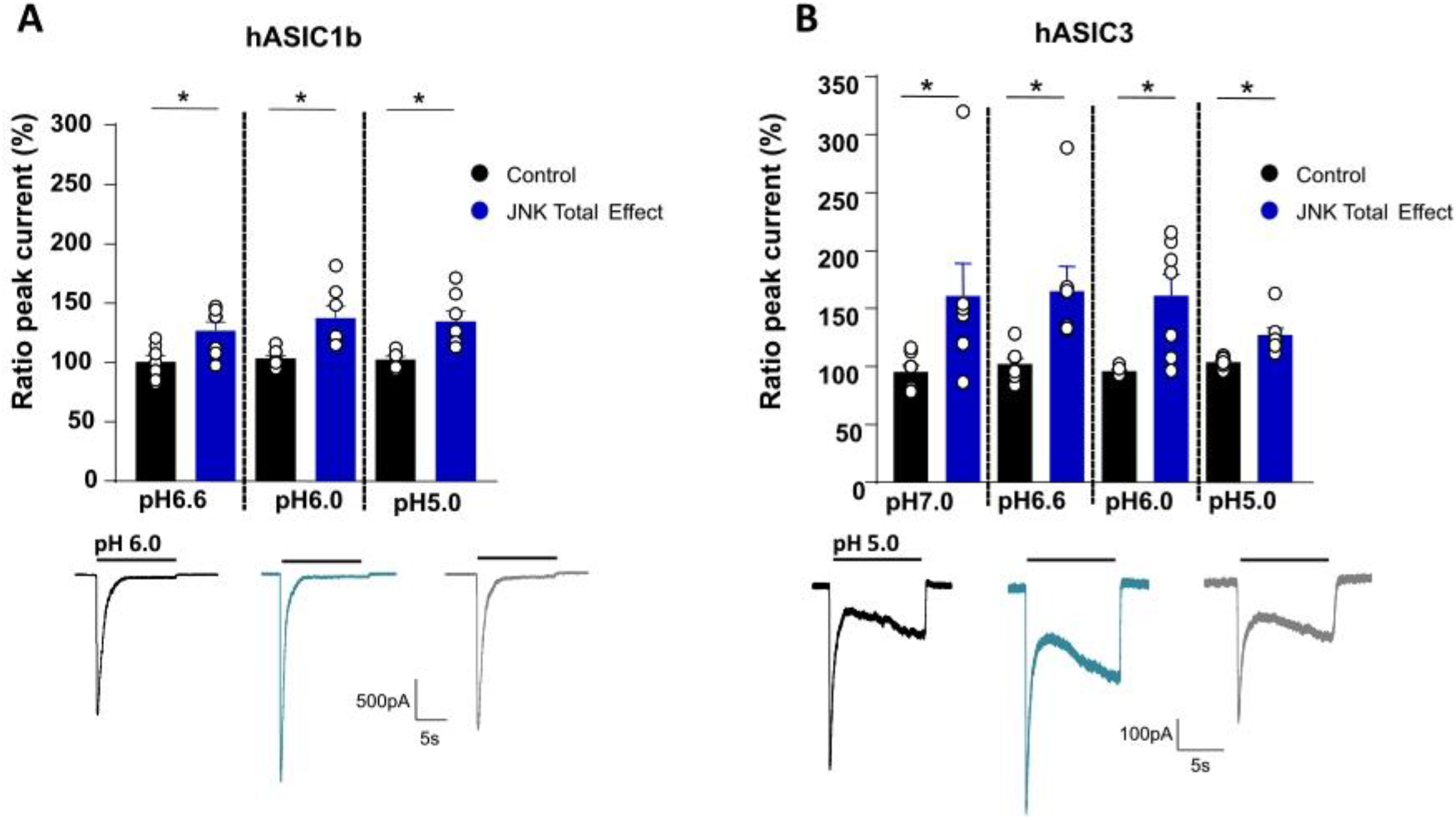
JNK-regulation of human ASIC1b and ASIC3 peak currents. **A,B**, Top: Potentiation by JNK (total effects) of recombinant H^+^-activated human ASIC1b (hASIC1b, A) and ASIC3 (hASIC3, B) peak currents expressed in HEK293 cells, activated at different pH (conditioning pH7.4) and recorded at −60mV (calculated and plotted as in Fig. 1B). n=7 cells for each channel subtype. Wilcoxon paired test versus control, * p<0.05. Bottom: Representative current traces in control condition (black), and after anisomycin (cyan) or SP600125 (grey) application.

These data show that short-term JNK activation exerts a positive regulation of the activity of recombinant rat and human ASIC1b- and ASIC3-containing channels, and that the presence of these subunits is necessary and sufficient to confer regulation to heteromeric channels combining subunits that are not *per se* regulated, like ASIC1a and ASIC2a.

### The JNK regulation is species-dependent and depends on a putative phosphorylation site within the intracellular domains of ASIC1b and ASIC3 subunits

To address the mechanism underlying the JNK-dependent potentiation of ASICs, we first performed a bioinformatic analysis on the intracellular segments of rASICs to identify putative phosphorylation sites. Interestingly, we identified a putative JNK2 phosphorylation site in the rASIC1b N-terminal domain (S59) as well as in the rASIC3 C-terminal domain (T512). These sites are conserved in rat, mouse and human except for mouse ASIC3 where the threonine is replaced by an alanine (Fig. 3A).

**Figure 3:**
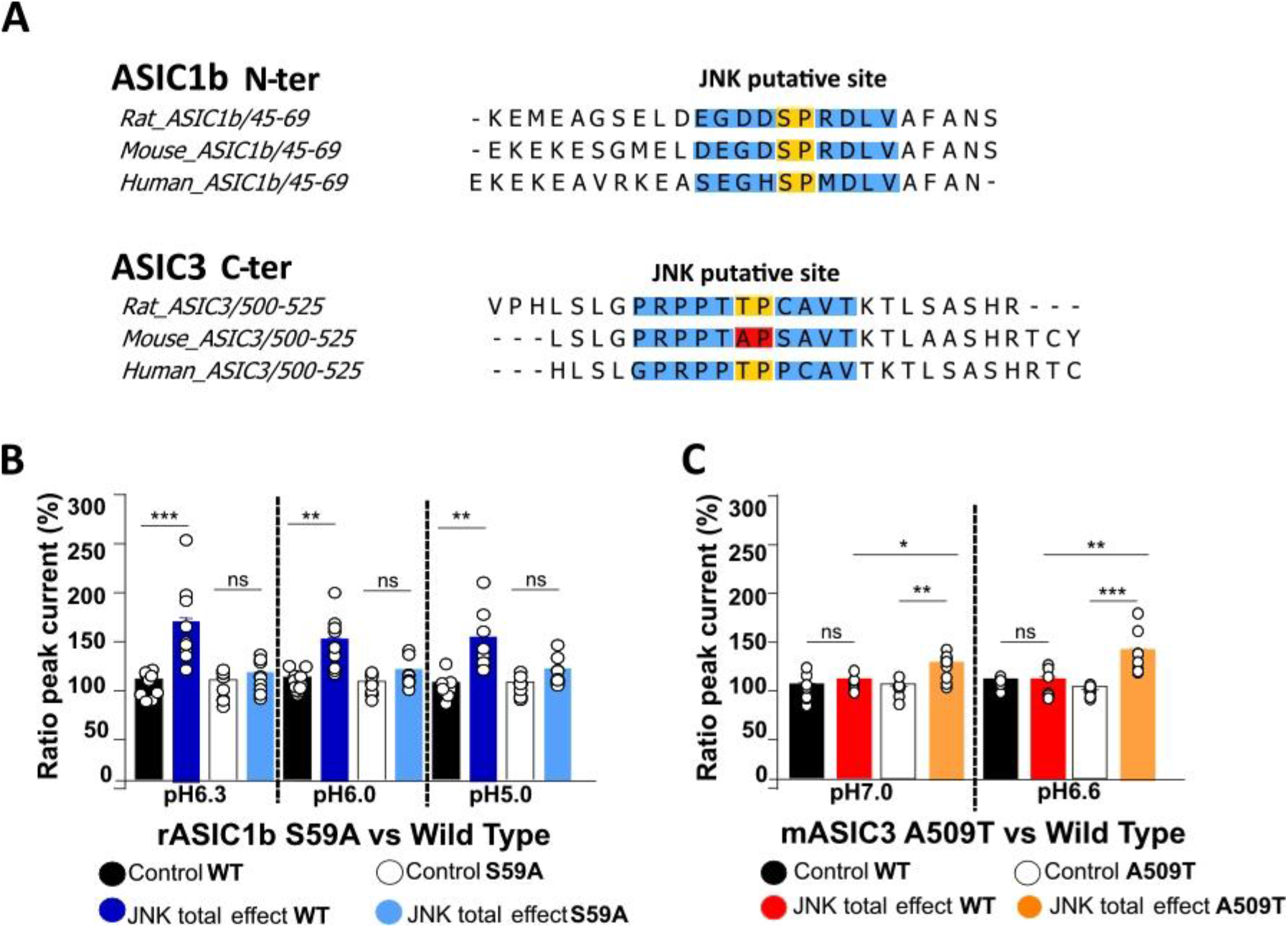
The JNK regulation is species-dependent and depends on a putative phosphorylation site within the intracellular domains of ASIC1b and ASIC3 subunits. **A,** Partial sequence alignments of rat, mouse and human ASIC1b and ASIC3 intracellular N- and C-terminal domain, respectively. The potential JNK phosphorylation sites found with GPS and ELM softwares are highlighted (phosphorylated S/T residues followed by P in blue). **B,C,** Potentiation by JNK (total effects) of H^+^-activated rASIC1b WT and S59A mutant (B), and of H^+^-activated mASIC3 WT and A509T mutant (C), recorded at different pH (calculated and plotted as in Fig. 1B). In B, one-way ANOVA followed by a Tukey post-hoc test (F(2, 26)=13.79, p<0.0001, n=9-12 cells). In C, one-way ANOVA followed by a Tukey post-hoc test (F(3, 28)=8.00, p=0.0005, n=6-9 cells). ***, p<0.001, **, p<0.005, *, p<0.05.

We next performed point mutations of the putative phosphorylation sites to test their possible involvement in the effects observed. The rASIC1b S59A mutant had biophysical properties similar to the wild-type channel (pH_0.5_ for activation and inactivation, inactivation time constant; not shown), but lost the regulation by JNK whatever the test pH used (Fig. 3B). Consistent with the lack of putative JNK phosphorylation site (Fig. 3A), the mouse ASIC3 channel failed to be potentiated by JNK (Fig 3C), but the mASIC3 A509T mutant in which the alanine is replaced by a threonine to create a putative JNK phosphorylation site, was significantly potentiated by the kinase (Fig 3C). Interestingly, since mouse and rat ASIC3 have more than 96% protein identity, mASIC3 A509 could be indeed considered as a “natural” mutation of the rat ASIC3 channel. Together with the A509T mutation that recreates in mouse the rat putative JNK phosphorylation site, this confirms the importance of this phosphorylation site in the JNK-dependent regulation of rat ASIC3.

These results suggest that potentiation of the ASIC1b- and ASIC3-containing channel activity by JNK is supported by a phosphorylation of the ASIC1b and ASIC3 subunits at intracellular S59 and T512, respectively. The importance of this later site is further confirmed by the fact that a single mutation of the putative phosphorylation residue is sufficient to confer JNK-regulation to mouse ASIC3 that is naturally insensitive.

### Native ASIC channels in DRG neurons are positively regulated by short-term JNK activation

The effects of JNK on native proton-gated currents in cultured DRG sensory neurons were then investigated. Rapid extracellular acidification from pH7.4 to pH5.0 on cultured mouse and rat DRG neurons hold at −80mV and randomly selected produced inward currents in all the neurons tested (n=101 and 49, respectively), with various shapes and kinetics. Neurons expressing transient ASIC-like currents, associated or not with a sustained current of variable amplitude (Supplementary Fig. 2A, traces #1 and #2) were selected, while neurons showing only a sustained current without any peak (Supplementary Fig. 2A, traces #3 and #4) were excluded. DRG neurons expressed a number of different, and often heteromeric, ASIC channels (Benson et al., 2002; Papalampropoulou-Tsiridou et al., 2020). We used a set of specific and potent ASIC-blocking toxins (Baron et al., 2013) to perform a pharmacological profiling of ASIC-like transient currents recorded in cultured DRG neurons to confirm the channel subtypes possibly associated. PcTx1, APETx2 and mambalgin-1 were indeed shown to block ASIC1a-containing (Escoubas et al., 2000), ASIC3-containing (Diochot et al., 2004), and both ASIC1a- and ASIC1b-containing (*i.e.*, ASIC1-containing) (Diochot et al., 2012) channels, respectively. An inhibition by at least 20% of the ASIC-like peak current by APETx2 (3μM), PcTx1 (20nM), or mambalgin-1 (1μM) was interpreted as a significant participation of ASIC3-containing channels, ASIC1a channels, or ASIC1-containing channels, respectively. The different pharmacological profiles of ASIC currents in neurons and the inferred ASIC channel subtypes possibly associated are listed in Table 1. Neurons expressing an ASIC-like current can be found into both small and medium/large diameter DRG populations (membrane capacitance <40pF and >40pF, respectively) whatever the display of a prominent shoulder on the falling phase of the action potential (not shown), a feature of nociceptive neurons (Petruska et al., 2000), consistent with the broad distribution of ASICs in the global sensory neuron population.

**Table 1:** Pharmacological profiles of DRG-expressed JNK-sensitive and -insensitive ASIC currents and their possible association with ASIC channel subtypes inferred from peptide toxin blocking selectivity. **Left,** number of sensory neurons expressing toxin-inhibited ASIC currents in mouse and rat primary cultures (JNK-sensitive versus total JNK-tested) and their distribution according to the different pharmacological profiles. JNK-dependent and -independent TNFα up-regulation of ASIC currents in acutely dissociated mouse DRG neurons is also shown. **Right,** ASIC channel subtypes possibly associated (+) or not (-) with the different pharmacological profiles determined by testing the inhibitory effects of mambalgin-1 (1μM), APETx2 (3 μM) and PcTx1 (20 nM). Channel subtypes regulated by JNK *in vitro* are indicated in bold. Only ASIC1b-containing channels are regulated in mouse, including heteromeric channels with ASIC3. Note that when different pharmacologically distinct types of channels are present together into a neuron (*e.g.*, ASIC1b-containing and ASIC3-containing channels), the proportion of each channel type can significantly vary depending on the percentage of inhibition induced by each toxin that can vary from 20% (threshold for inhibition) to 100%.

| Number of neurons |  |  | Toxin selectivity | mambalgin-1 |  | APETx2 |
| --- | --- | --- | --- | --- | --- | --- |
| JNK-sensitive/total | | TNFα-sensitive/JNK-sensitive/total | | PcTx1 | | |
| Mouse (32/40) | Rat (18/20) | Mouse TNFα (11/9/19) | DRG currents blocked by : | ASIC1a | ASIC1a/2a or ASIC1b-containing | ASIC3-containing |
| 5/9 | 6/6 | 2/1/7 | mambalgin-1 + APETx2 + PcTx1 | + | + | + |
| 17/21 | 5/5 | 6/6/7 | mambalgin-1 + APETx2 | - | + | + |
|  | 3/5 |  | mambalgin-1 + PcTx1 | + | + | - |
| 10/10 |  | 3/2/5 | mambalgin-1 | - | + | - |
|  | 4/4 |  | APETx2 | - | - | + |

The MAPK activator anisomycin was then applied on mouse and rat ASIC-expressing neurons (n= 40 and 20, respectively) and showed little potentiating effect on ASIC peak current associated with various pharmacological profiles (Fig. 4A-C), whereas subsequent perfusion of the JNK inhibitor SP600125 induced a marked reduction of the current amplitude below the control level, thus revealing a high basal level of regulation of ASIC currents by JNK in at least 80% of the neurons tested in mouse and rat (32/40 and 18/20 neurons, respectively, Table 1). The culture process used is indeed known to increase the basal activity of the JNK pathway (Kristiansen and Edvinsson, 2010). A total JNK effect of +100.8±17.4% at pH5.0 and +107±19.0% at pH6.6 in mouse DRG neurons, and of +78.8±13.6% at pH5.0 and +66.6±15.5% at pH6.6 in rat DRG neurons was calculated (Fig. 4A-C). Interestingly, a small subset of mouse neurons expressed APETx2-sensitive (*i.e.*, ASIC3-containing channels) but JNK-insensitive currents (Table 1 and Fig. 4D, upper traces) while all currents sensitive to APETx2 were sensitive to JNK in rat (Table 1 and Fig. 4D, lower trace). This is consistent with the previous experiments in HEK293 cells (Figs. 1, 2 and 3) and with the fact that the JNK regulation in mice solely involves the ASIC1b, but not the ASIC3, subunit. A large proportion of neurons expressing APETx2-sensitive currents in mouse were however regulated by JNK (Table 1 and Fig. 4A, lower traces), suggesting that they flow through heteromeric ASIC3-containing channels also including the JNK-sensitive ASIC1b subunit. Two rat neurons displaying currents sensitive to mambalgin-1 and PcTx1, thus likely expressing a majority of ASIC1a channels, were not sensitive to JNK (Table 1 and Fig. 4D, middle-down traces), but neurons likely expressing ASIC1b-containing channels (*i.e.,* sensitive to the toxin mambalgin-1 but not PcTx1) all showed JNK-regulated currents (Table 1 and Fig. 4D, middle-up traces), in good agreement with our previous experiments on recombinant ASIC channels in HEK293 cells.

**Figure 4:**
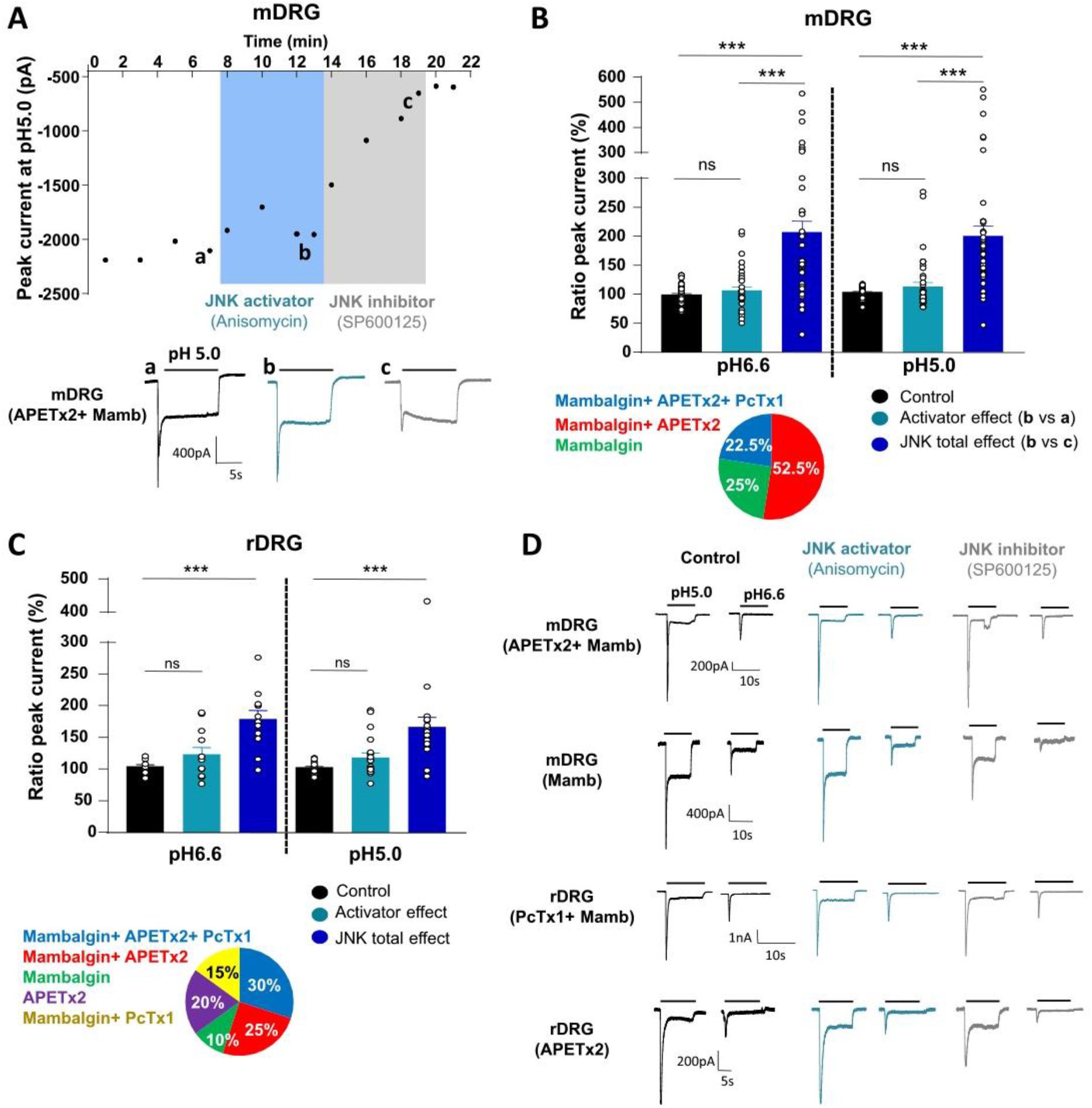
JNK regulates native ASIC peak currents in cultured rodent DRG neurons. **A**, Top: Representative time course of mouse DRG (mDRG) neuron ASIC peak current amplitude recorded at −80mV upon stimulation by a shift from resting pH7.4 to pH5.0, and extracellular perfusion with anisomycin (50μM) and the JNK inhibitor SP600125 (50μM). Bottom: Representative current traces of APETx2 and mambalgin-1-sensitive currents (see Table 1 for details). **B,** Potentiation by JNK of H^+^-activated mouse DRG (mDRG) neuron ASIC peak current recorded at different pH (calculated and plotted as in Fig. 1B), n=36-40. One-way ANOVA (F(2,78)=22.04 p<0.0001) followed by Tukey post-hoc test. ***, p<0.0001. Bottom: Pie chart showing the proportion of ASIC peak currents with different pharmacological profiles tested for JNK regulation in mouse DRG neurons (n=40) and determined by the effects of the three ASIC inhibitory toxins mambalgin-1 (1μM), APETx2 (3 μM) and PcTx1 (20 nM). A 20% threshold for inhibition has been used. See Table 1 for details and interpretation. **C**, Potentiation by JNK of H^+^-activated rat DRG (rDRG) neuron ASIC peak current recorded at different pH (same protocol as in B), n=12-20. One-way ANOVA (F(2,38)=10.62, p=0.0002) followed by Tukey post-hoc test ***, p<0.0001. Bottom: Pharmacological profiles (determined as in B, see Table 1 for interpretations) of the ASIC peak currents in rat DRG neurons tested for JNK regulation (n=20). **D**, Representative current traces of rat and mouse DRG neuron ASIC currents with different pharmacological profiles regarding inhibition by mambalgin-1 (Mamb), PcTx1 and APETx2 in control condition (black), and after anisomycin and SP600125 application (cyan and grey, respectively). The toxins inhibiting the current by at least 20% were indicated in brackets. Currents were recorded at −80mV and activated at two different pHs. See Table 1 for interpretation of the pharmacological profiles.

All together, these data show that the activity of native ASIC1b- and ASIC3-containing channels is up-regulated by short-term JNK activation in most mouse and rat DRG neurons.

### JNK-dependent up-regulation of native ASIC currents in DRG neurons by the proinflammatory cytokine TNFα

The ability of endogenous pain-related mediators to regulate native ASIC channels in DRG neurons *via* the JNK-pathway was then tested. The effect of TNFα, a pain-related cytokine and a well-known activator of receptor-mediated JNK activity (Natoli et al., 1997), was tested in acutely dissociated mouse DRG neurons (3-10h in culture) deprived of neurotrophic factors in order to limit basal activation of JNK. The ASIC-like peak currents were characterized in these neurons as previously described and displayed pharmacological profiles close to those established in primary cultures (Fig. 5B versus Fig. 4B, lower panels). The subsequent perfusion of TNFα (10ng/ml) on these neurons led to a rapid (within 5 minutes) increase of the ASIC peak current amplitude in more than half of the neurons tested (11/19) (Fig. 5A,B; Table 1). Subsequent perfusion of the JNK inhibitor SP600125 induced a decrease of the current amplitude in 9/11 TNFα-responding neurons, including 8 neurons that are only sensitive to mambalgin-1 and not to PcTx1, *i.e.*, likely expressing ASIC1b-containing channels (Table 1). There was no major difference in the magnitude of increase induced by TNFα (TNFα effect compared to control) and the TNFα-JNK total effect (+36.4±11.1% *vs* +58.1±11.8% at pH5.0, and +57.2±18.4 *vs* +62.3±15.8 at pH6.6, not statistically significantly different, Fig. 5B), showing no significant basal activation of the JNK pathway in the absence of neurotrophic factors. The TNFα-induced increase of ASIC peak current amplitude therefore mainly involved activation of the JNK pathway. However, the potentiating effect of TNFα was not reversed by SP600125 in two neurons out of 11, including a neuron that predominantly expressed ASIC1b-containing channels (*i.e.*, sensitive to mambalgin-1 and not to PcTx1) (Table 1), suggesting a possible minor involvement of other JNK-independent TNFα-associated regulation(s) of ASICs in DRG neurons.

**Figure 5:**
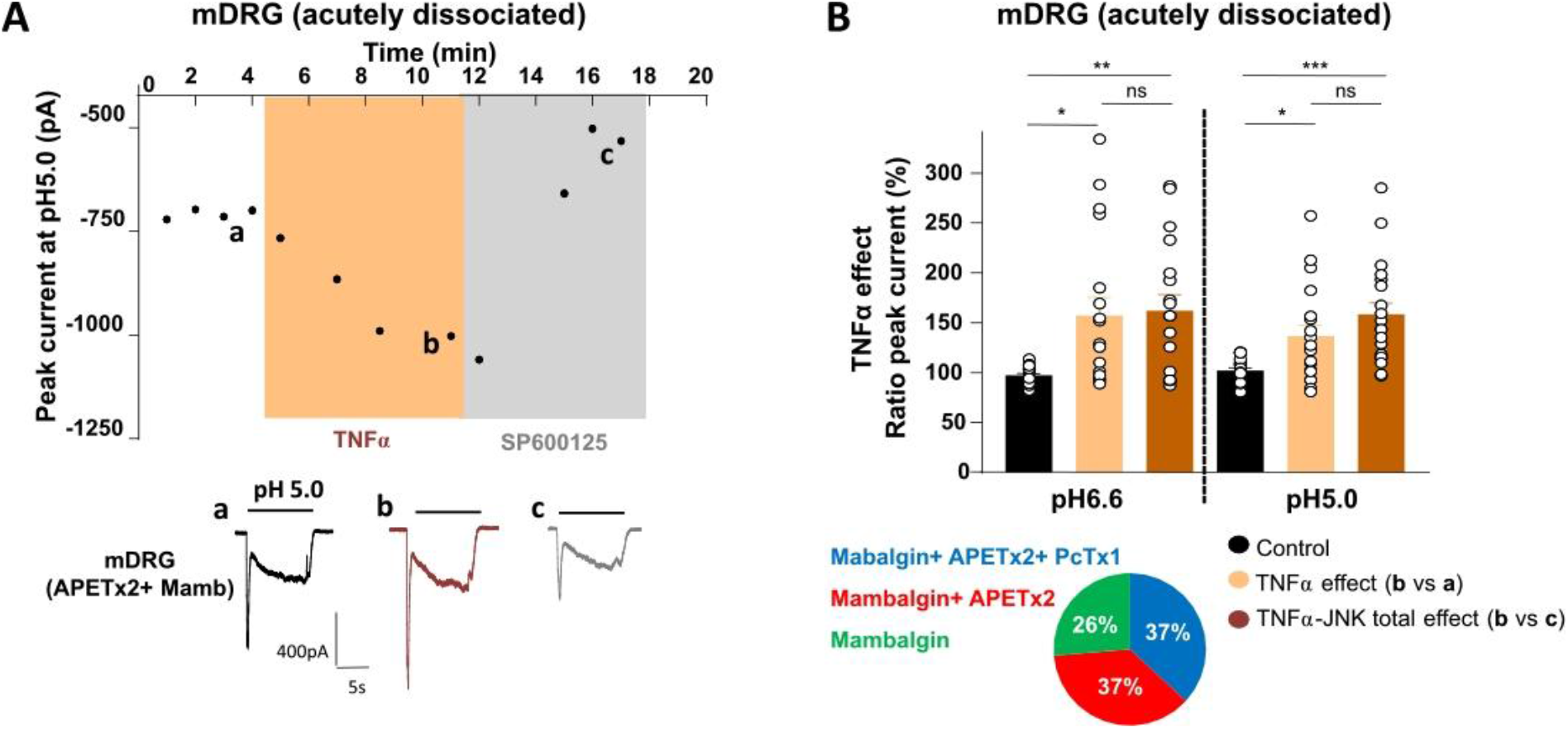
TNFα potentiates H^+^-activated ASIC peak currents from acutely dissociated mouse sensory neurons through JNK activation. **A**, Top: Representative time course of mouse DRG (mDRG) ASIC peak current amplitude upon stimulation at pH5.0 from conditioning pH7.4 and extracellular perfusion with TNFα (10ng/ml) and the JNK inhibitor SP600125 (50μM). Bottom: Representative pH5.0-evoked mambalgin-1- (Mamb) and APETx2-sensitive current traces (see Table 1). **B**, Mean potentiation by TNFα of H^+^-activated ASIC peak currents in mouse DRG (mDRG) neurons recorded at different pH (ratio b/a, plotted as in Fig. 1B) and comparison with the TNFα-JNK total effect (ratio b/c). n=18-19, One-way ANOVA (F(2,36)=10.47, p=0.0004) followed by Tukey post-hoc test, ***, p<0.001, **, p<0.01, *, p<0.05. Bottom: Pharmacological profiles of the ASIC peak currents in mouse DRG neurons tested for TNFα regulation (as determined in Fig.4B; n=19).

The proinflammatory cytokine TNFα therefore appears as a fast and potent up-regulator of ASIC1b-containing channel activity in mouse sensory neurons mainly through the activation of the JNK pathway.

### Potentiation by JNK of the acid-induced cutaneous pain in inflammatory conditions in mice and implication of peripheral ASIC1-containing channels

The physiopathological relevance of JNK regulation of ASIC channels was tested *in vivo* on the spontaneous pain behavior induced by subcutaneous intraplantar (*i.pl.*) injection of a pH5.0 solution in the mouse hindpaw. A pharmacological approach has been used where compounds (or corresponding vehicles) were pre-injected ten minutes before their co-injection with the pH5.0 solution. In naïve mice, pH5.0 *i.pl.* injection (preceded by pre-injection of vehicle) induced a very small behavioral response during the first 5 minute-interval (5.9 ± 1.2 s, n=26, Fig. 6A). In the presence of anisomycin (500 μM), the acid-induced response was significantly increased (10.8 ± 2.9 s, n=23), and was also stronger than the effect of anisomycin alone (6.3 ± 2.3 s, n=23, Fig. 6A). Although these results already show some potentiation by anisomycin of the pH5.0-induced pain in naïve animals, the small amplitude of the effects precluded further investigation in these conditions.

**Figure 6:**
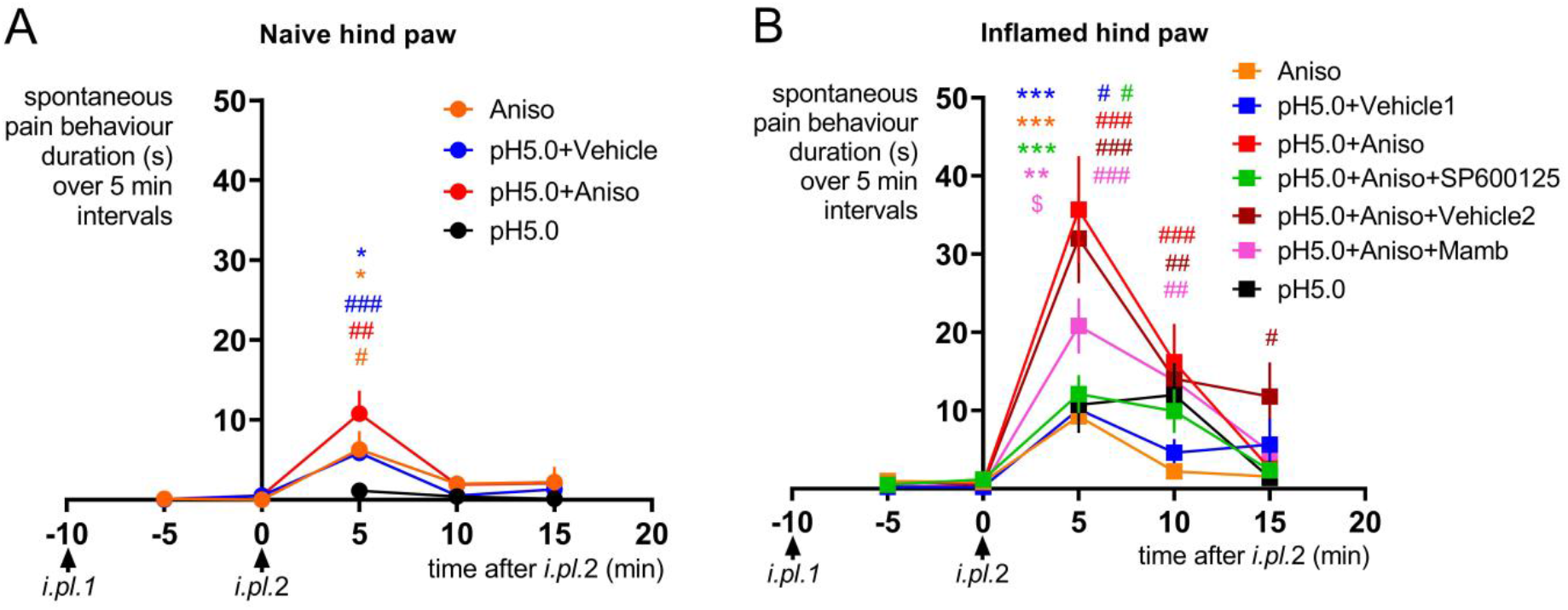
ASIC1-containing channels contribute to the JNK-induced potentiation of cutaneous acid-triggered pain in mice. **A**, Kinetics of spontaneous pain behavior duration (s) measured over 5 min-intervals in naïve mice hindpaw. Single *i.pl.* injection of pH5.0 (black symbols, n=28) and double *i.pl.* injection with drugs being pre-injected alone (*i.pl.* 1) 10 min before their co-injection with pH5.0 (*i.pl.2):* Anisomycin alone (500 μM, Aniso, orange symbols, n=23), pH5.0 + vehicle (DMSO 0.5%, blue symbols, n= 26), pH5.0+Anisomycin (500 μM, red symbols, n=23). **B**, Kinetics of spontaneous pain behavior duration (s) measured over 5 min-intervals in mice with local hindpaw inflammation (carrageenan 2%, 3 hours). Single *i.pl.* injection of pH5.0 (black symbols, n=25) and double *i.pl.* injection protocol as described in A: Anisomycin alone (500 μM, Aniso, orange symbols, n=25), pH5.0+vehicle1 (DMSO 0.5%, blue symbols, n= 26), pH5.0+Anisomycin (500 μM, red symbols, n=25), pH5.0+Anisomycin+SP600125 (500μM each, green symbols, n=25), pH5.0+Anisomycin+vehicle2 (500μM and 0.05% BSA, respectively, dark red symbols, n=25), pH5.0+Anisomycin+Mambalgin-1 (500 μM and 34 μM, respectively, pink symbols, n=23). Data are presented as mean ± SEM as a function of time. For data obtained with the double *i.pl.* protocol, # p<0.05, ## p<0.01, ### p<0.005 compared to the 5 min interval before *i.pl.*2 injection with a two-way Anova followed by a Dunnett post-hoc paired comparison on the same mice. * p<0.05, ** p<0.01, *** p<0.005 compared with pH5.0+Anisomycin with a two-way Anova followed by a Dunnett post-hoc test. $ p<0.05, compared with pH5.0+Anisomycin+vehicle2 with a two-way Anova followed by a Dunnett post-hoc test.

A possible potentiation of acid-induced cutaneous pain by JNK was also tested in inflammatory conditions after an *i.pl.* injection of carrageenan in the hindpaw 3 hours before testing the acid pH5.0-induced spontaneous pain. The pH5.0 injection induced a 10.7 ± 3.6 s (n=25) mean response during the first 5 minute-interval (Fig. 6B), with 76% of the tested mice showing a response longer than 3 s compared to only 14% of naïve mice, thus showing that the local hindpaw inflammation not only increased the duration of the response but above all the number of responding mice. Anisomycin induced a significantly enhanced pH5.0-induced pain behavior during the first 5 minutes (35.7 ± 6.8 s, n=25) compared to the effect of vehicle (10.1 ± 2.3 s, n=26, Fig. 6B) or to the effect of anisomycin alone (9.3 ± 2.1 s, n=25). The potentiating effect of anisomycin on acidic pain was reversed by the JNK inhibitor SP600125 (500 μM), leading to a behavioral response of 12.1 ± 2.4 s (n=25, Fig. 6B) that was similar to the response in the absence of anisomycin. JNK has therefore a major contribution to the anisomycin-induced acidic pain potentiation in inflammatory conditions. The response was significantly reduced in the presence of mambalgin-1 (34 μM), the specific ASIC1-containing channel inhibitor, to 20.8 ± 3.5 s (n=23) during the first 5 minutes compared to 32.0 ± 5.7 s (n= 25) in the presence of the corresponding vehicle (Fig. 6B).

These data show that ASIC1-containing channels are involved in the JNK-potentiated, acid-induced cutaneous pain *in vivo* in inflammatory conditions.

## Discussion

By combining electrophysiological approaches with specific pharmacologic tools, we identified a new JNK-mediated fast potentiation of the activity of ASIC1b- and ASIC3-containing channels. This regulation may involve a direct phosphorylation of the intracellular domains of the ASIC subunits since i) it is disrupted by mutation of a putative JNK phosphorylation site in the ASIC1b N-terminal domain (S59A), ii) it is absent in mouse ASIC3 naturally lacking the phosphorylation site because of the presence of an alanine (A509) instead of a threonine, but can be acquired by introducing a point mutation (A509T) recreating a JNK phosphorylation site. Native ASIC channels in DRG neurons are potentiated by JNK with a functional profile consistent with the data found from recombinant channels in HEK293 cells, and this JNK signaling pathway is involved in the fast potentiation of native ASIC currents by the proinflammatory cytokine TNFα in sensory neurons. Finally, ASIC1-containing channels contribute to the JNK potentiating effect on cutaneous acid-triggered pain in inflammatory conditions in mice.

### MAPKs as potent post-translational regulators of ASIC channels

Our data support a role for JNK in the post-translational regulation of ASIC channels. We have combined in our protocols an activator (anisomycin) and an inhibitor (SP600125) that displays at the concentration used (50μM) a good selectivity for JNK versus the other MAPKs (Bain et al., 2007). In addition, the ASIC current potentiation strictly depends on putative JNK phosphorylation sites in ASIC1b and ASIC3 subunits. Even if SP600125 has the potency to inhibit to some extent other kinases (*e.g.*, DYRK, PDK1, CK2 or SGK1) (Bain et al., 2007), the predictive phosphorylation sites for theses kinases present in the ASIC1b and/or ASIC3 intracellular domains are clearly different from the two that has been shown to be mandatory for the effect described here. In addition, some of these predictive phosphorylation sites such as SGK1 are also present in the ASIC1a or ASIC2a subunits, which are not affected by SP600125. A role for JNK is therefore strongly supported by these complementary elements, even if minor contributions of other signaling pathways cannot be completely excluded.

A large range of mediators of various origins have been shown to directly or indirectly modulate the activity of ASIC channels. Several pain-related signaling pathway targeting ASICs have been previously characterized, including PKC (Baron et al., 2002), PKA (Leonard et al., 2003) or phosphoinositide 3-kinase (PI3K)-protein kinase B (PKB/Akt) (Duan et al., 2012). A MAPK-dependent transcriptional regulation of ASIC3 has been involved in the effect of NGF (Mamet et al., 2003; Chaumette et al., 2020) or interleukin 1β (Ross et al., 2016), and a MAPK-dependent transcriptional regulation of ASIC1a has also been associated to the effect of interleukin 6 (Zhou et al., 2015) or oxidative stress (Wu et al., 2020), but a MAPK-dependent phosphorylation of ASIC channels has never been described so far. Our data are consistent with a phosphorylation by JNK of the ASIC1b and ASIC3 (except mouse ASIC3) subunits. The time scale of the effect (less than 10 minutes) makes an indirect transcriptional contribution to the effect unlikely. JNK may therefore participate in nociceptive neurons to the generation of pain hypersensitivity through transcription-independent, direct phosphorylation of ion channels besides its well-known transcription-dependent modulation of pain-related genes. The biophysical properties of ASIC1b channels including gating, proton sensitivity, inactivation time constant and conductance, are not affected by the JNK regulation, which may suggest an increase in the opening probability of the channel or an effect mediated at least in part through an increase in channel trafficking to the plasma membrane. Enhanced forward trafficking and increased surface expression of ASICs *via* direct phosphorylation has been already reported for the regulation of ASIC1a by BDNF through the PI3-kinase/Akt pathway in the spinal cord (Duan et al., 2012). The effect of JNK phosphorylation on a given target is often linked to the establishment of new protein-protein interactions (Zeke et al., 2016). This phosphorylation could therefore be a switch to increase channel number at the plasma membrane, as shown previously (Simonetti et al., 2014), or to prevent ion channels to be internalized.

### Native proton-gated currents of sensory neurons are strongly potentiated by JNK to modulate pain sensing

Our results demonstrate that the vast majority of DRG neurons expressing ASIC currents (including small and medium/large neurons) are positively regulated by JNK signaling. The molecular identity of these currents can be, at least partially, inferred from their biophysical properties and their sensitivity to different ASIC toxins specifically blocking complementary set of homomeric and/or heteromeric channels (Baron et al., 2013). Some JNK-insensitive currents have been only recorded in ASIC1a-expressing neurons in rat, or in ASIC3-expressing neurons in mouse (all ASIC3-expressing neurons display JNK-regulated ASIC currents in rat). It fits with our experiments on recombinant channels in HEK293 cells showing a regulation supported by both the ASIC1b and ASIC3 subunits in rat but only the ASIC1b subunit in mouse. However, more than 70% of mouse ASIC3-expressing neurons display JNK-regulated currents despite the lack of regulation of ASIC3 subunit, suggesting the presence in DRG neurons of our primary cultures of a high proportion of ASIC3 heteromeric channels also containing ASIC1b. Multiplexed *in situ* hybridization has confirmed a significant co-localization of ASIC1b and ASIC3 transcripts in mouse DRG neurons (Chang et al., 2019; Papalampropoulou-Tsiridou et al., 2020). It further supports, together with the high proportion of JNK-regulated ASIC1b-expressing neurons in mouse, a broad functional expression of ASIC1b in sensory neurons. This is fully consistent with our previous work indicating a significant role for sensory-neuron expressed ASIC1b-containing channels in pain (Diochot et al., 2012; Diochot et al., 2016). Our *in vivo* experiments based on local application of a specific ASIC blocker, further support the involvement of peripheral ASIC1-containing channels, most probably ASIC1b-containing channels regarding the *in vitro* data on recombinant and native ASICs, in the JNK-potentiated acidic pain in inflammatory conditions. Interestingly, the strong potentiating effect observed in inflammatory conditions is consistent with a role for ASIC1b in producing “hyperalgesic priming”, a process that has been proposed to be involved in the transition from acute to chronic pain, as recently suggested in a preclinical model of muscle pain (Chang et al., 2019). The effect was however only partially inhibited by the specific ASIC blocker, suggesting the involvement, at least in mice, of other JNK-regulated and ASIC-independent processes.

Direct regulation of DRG-specific ASIC1b- and ASIC3-containing channels by JNK represents a new mechanism for pathological potentiation of ASIC channels upon activation of this signaling pathway in peripheral sensory neurons. Extracellular mediators contributing to pain sensitization can participate in this process like the cytokine TNFα (Zelenka et al., 2005) that rapidly increases the amplitude of ASIC currents mainly in a JNK-dependent manner as shown here for the first time. However, another study reported a lack of potentiating effect of TNFα on rat vagal sensory neuron rapid transient current mediated by ASICs but a significant increase of the slow sustained current mediated by TRPV1 through a COX2-dependant mechanism (Hsu et al., 2017). Differences in the experimental conditions (origin of the neurons and culture conditions, TNFα concentration, perfusion time) could explain these different results. Interestingly, the JNK-dependent regulation of the activity of ASIC1b- and ASIC3-containing channels in sensory neurons is reminiscent to the one previously mentioned for BDNF, PI3-kinase/Akt pathway and ASIC1a channels in the spinal cord (Duan et al., 2012).

In conclusion, our results identify a novel mechanism of post-translational regulation of sensory neuron-expressed ASIC1b- and ASIC3-containing channels by JNK, probably *via* direct phosphorylation of the channels suggesting that ion channels can be direct downstream effectors of JNK in nociceptors. Our *in vivo* results on cutaneous pain illustrate the important role this regulation may have in various pathophysiological pain conditions involving the JNK signaling pathway including inflammatory, neuropathic and migraine pain where ASIC1b and/or ASIC3-containing channels have been involved (Deval et al., 2008; Diochot et al., 2012; Yan et al., 2013; Diochot et al., 2016; Verkest et al., 2018). In addition, we provide further evidence of a large functional expression of ASIC1b-containing channels in sensory neurons, supporting the emerging role in pain of this ASIC isoform. The JNK regulation is conserved in human ASIC1b and ASIC3 channels and interfering with this regulation at the channel level might be of potential therapeutic benefit against pain.

## Author contributions

C.V., E.L. and A.B. designed research; C.V., S.D. and A.B. performed research; all the authors analyzed data and wrote the paper.

## Acknowledgements

We thank L. Meneux, E. Deval, J. Noël, M. Salinas, A. Negm, M. Chafai, L. Pidoux, K. Delanoë and B. Labrum for helpful discussions, V. Friend and J. Salvi-Leyral for expert technical assistance, and V. Berthieux for secretarial assistance. This work was supported by the Centre National de la Recherche Scientifique, the Institut National de la Santé et de la Recherche Médicale, and the Agence Nationale de la Recherche (ANR-17-CE18-0019, ANR-17-CE16-0018 and ANR-11-LABX-0015-01).

## References

Askwith CC, Cheng C, Ikuma M, Benson C, Price MP, Welsh MJ (2000) Neuropeptide FF and FMRFamide potentiate acid-evoked currents from sensory neurons and proton-gated DEG/ENaC channels. Neuron 26:133–141.

Bain J, Plater L, Elliott M, Shpiro N, Hastie CJ, McLauchlan H, Klevernic I, Arthur JSC, Alessi DR, Cohen P (2007) The selectivity of protein kinase inhibitors: a further update. The Biochemical Journal 408:297–315.

Baron A, Deval E, Salinas M, Lingueglia E, Voilley N, Lazdunski M (2002) Protein kinase C stimulates the acid-sensing ion channel ASIC2a via the PDZ domain-containing protein PICK1. The Journal of Biological Chemistry 277:50463–50468.

Baron A, Diochot S, Salinas M, Deval E, Noël J, Lingueglia E (2013) Venom toxins in the exploration of molecular, physiological and pathophysiological functions of acid-sensing ion channels. Toxicon: Official Journal of the International Society on Toxinology 75:187–204.

Benson CJ, Xie J, Wemmie JA, Price MP, Henss JM, Welsh MJ, Snyder PM (2002) Heteromultimers of DEG/ENaC subunits form H+-gated channels in mouse sensory neurons. Proceedings of the National Academy of Sciences of the United States of America 99:2338–2343.

Bohlen CJ, Chesler AT, Sharif-Naeini R, Medzihradszky KF, Zhou S, King D, Sánchez EE, Burlingame AL, Basbaum AI, Julius D (2011) A heteromeric Texas coral snake toxin targets acid-sensing ion channels to produce pain. Nature 479:410–414.

Cadiou H, Studer M, Jones NG, Smith ESJ, Ballard A, McMahon SB, McNaughton PA (2007) Modulation of acid-sensing ion channel activity by nitric oxide. The Journal of Neuroscience: The Official Journal of the Society for Neuroscience 27:13251–13260.

Chang C-T, Fong SW, Lee C-H, Chuang Y-C, Lin S-H, Chen C-C (2019) Involvement of Acid-Sensing Ion Channel 1b in the Development of Acid-Induced Chronic Muscle Pain. Frontiers in Neuroscience 13:1247.

Chaumette T, Delay L, Barbier J, Boudieu L, Aissouni Y, Meleine M, Lashermes A, Legha W, Antraigue S, Carvalho FA, Eschalier A, Ardid D, Moqrich A, Marchand F (2020) c-Jun/p38MAPK/ASIC3 pathways specifically activated by NGF through TrkA is crucial for mechanical allodynia development. Pain 161:1109–1123.

Deval E, Lingueglia E (2015) Acid-Sensing Ion Channels and nociception in the peripheral and central nervous systems. Neuropharmacology 94:49–57.

Deval E, Noël J, Lay N, Alloui A, Diochot S, Friend V, Jodar M, Lazdunski M, Lingueglia E (2008) ASIC3, a sensor of acidic and primary inflammatory pain. The EMBO journal 27:3047–3055.

Dinkel H et al. (2012) ELM--the database of eukaryotic linear motifs. Nucleic Acids Research 40:D242–251.

Diochot S, Baron A, Rash LD, Deval E, Escoubas P, Scarzello S, Salinas M, Lazdunski M (2004) A new sea anemone peptide, APETx2, inhibits ASIC3, a major acid-sensitive channel in sensory neurons. The EMBO journal 23:1516–1525.

Diochot S, Alloui A, Rodrigues P, Dauvois M, Friend V, Aissouni Y, Eschalier A, Lingueglia E, Baron A (2016) Analgesic effects of mambalgin peptide inhibitors of acid-sensing ion channels in inflammatory and neuropathic pain. Pain 157:552–559.

Diochot S, Baron A, Salinas M, Douguet D, Scarzello S, Dabert-Gay A-S, Debayle D, Friend V, Alloui A, Lazdunski M, Lingueglia E (2012) Black mamba venom peptides target acid-sensing ion channels to abolish pain. Nature 490:552–555.

Duan B, Liu D-S, Huang Y, Zeng W-Z, Wang X, Yu H, Zhu MX, Chen Z-Y, Xu T-L (2012) PI3-kinase/Akt pathway-regulated membrane insertion of acid-sensing ion channel 1a underlies BDNF-induced pain hypersensitivity. The Journal of Neuroscience: The Official Journal of the Society for Neuroscience 32:6351–6363.

Escoubas P, De Weille JR, Lecoq A, Diochot S, Waldmann R, Champigny G, Moinier D, Ménez A, Lazdunski M (2000) Isolation of a tarantula toxin specific for a class of proton-gated Na+ channels. The Journal of Biological Chemistry 275:25116–25121.

Francois A, Kerckhove N, Meleine M, Alloui A, Barrere C, Gelot A, Uebele VN, Renger JJ, Eschalier A, Ardid D, Bourinet E (2013) State-dependent properties of a new T-type calcium channel blocker enhance Ca(V)3.2 selectivity and support analgesic effects. Pain 154:283–293.

Hsu C-C, Lin YS, Lin R-L, Lee L-Y (2017) Immediate and delayed potentiating effects of tumor necrosis factor-alpha on TRPV1 sensitivity of rat vagal pulmonary sensory neurons. American Journal of Physiology Lung Cellular and Molecular Physiology 313:L293–L304.

Huang D, Ren L, Qiu C-S, Liu P, Peterson J, Yanagawa Y, Cao Y-Q (2016) Characterization of a mouse model of headache. Pain 157:1744–1760.

Hucho T, Levine JD (2007) Signaling Pathways in Sensitization: Toward a Nociceptor Cell Biology. Neuron 55:365–376.

Kristiansen KA, Edvinsson L (2010) Neurogenic inflammation: a study of rat trigeminal ganglion. The Journal of Headache and Pain 11:485–495.

Leonard AS, Yermolaieva O, Hruska-Hageman A, Askwith CC, Price MP, Wemmie JA, Welsh MJ (2003) cAMP-dependent protein kinase phosphorylation of the acid-sensing ion channel-1 regulates its binding to the protein interacting with C-kinase-1. Proceedings of the National Academy of Sciences of the United States of America 100:2029–2034.

Li W-G, Yu Y, Zhang Z-D, Cao H, Xu T-L (2010) ASIC3 channels integrate agmatine and multiple inflammatory signals through the nonproton ligand sensing domain. Molecular Pain 6:88.

Mamet J, Lazdunski M, Voilley N (2003) How nerve growth factor drives physiological and inflammatory expressions of acid-sensing ion channel 3 in sensory neurons. The Journal of Biological Chemistry 278:48907–48913.

Marra S, Ferru-Clement R, Breuil V, Delaunay A, Christin M, Friend V, Sebille S, Cognard C, Ferreira T, Roux C, Euller-Ziegler L, Noel J, Lingueglia E, Deval E (2016) Non-acidic activation of pain-related Acid-Sensing Ion Channel 3 by lipids. EMBO J 35:414–428.

Natoli G, Costanzo A, Ianni A, Templeton DJ, Woodgett JR, Balsano C, Levrero M (1997) Activation of SAPK/JNK by TNF Receptor 1 Through a Noncytotoxic TRAF2-Dependent Pathway. Science 275:200–203.

Obata K, Yamanaka H, Kobayashi K, Dai Y, Mizushima T, Katsura H, Fukuoka T, Tokunaga A, Noguchi K (2004) Role of mitogen-activated protein kinase activation in injured and intact primary afferent neurons for mechanical and heat hypersensitivity after spinal nerve ligation. The Journal of Neuroscience: The Official Journal of the Society for Neuroscience 24:10211–10222.

Papalampropoulou-Tsiridou M, Labrecque S, Godin AG, De Koninck Y, Wang F (2020) Differential Expression of Acid - Sensing Ion Channels in Mouse Primary Afferents in Naïve and Injured Conditions. Frontiers in Cellular Neuroscience 14:103.

Petho G, Reeh PW (2012) Sensory and signaling mechanisms of bradykinin, eicosanoids, platelet-activating factor, and nitric oxide in peripheral nociceptors. Physiological Reviews 92:1699–1775.

Petruska JC, Napaporn J, Johnson RD, Gu JG, Cooper BY (2000) Subclassified acutely dissociated cells of rat DRG: histochemistry and patterns of capsaicin-, proton-, and ATP-activated currents. Journal of neurophysiology 84:2365–2379.

Ross JL, Queme LF, Cohen ER, Green KJ, Lu P, Shank AT, An S, Hudgins RC, Jankowski MP (2016) Muscle IL1β Drives Ischemic Myalgia via ASIC3-Mediated Sensory Neuron Sensitization. The Journal of Neuroscience: The Official Journal of the Society for Neuroscience 36:6857–6871.

Salinas M, Lazdunski M, Lingueglia E (2009) Structural elements for the generation of sustained currents by the acid pain sensor ASIC3. The Journal of Biological Chemistry 284:31851–31859.

Simonetti M, Agarwal N, Stösser S, Bali KK, Karaulanov E, Kamble R, Pospisilova B, Kurejova M, Birchmeier W, Niehrs C, Heppenstall P, Kuner R (2014) Wnt-Fzd signaling sensitizes peripheral sensory neurons via distinct noncanonical pathways. Neuron 83:104–121.

Smith ES, Cadiou H, McNaughton PA (2007) Arachidonic acid potentiates acid-sensing ion channels in rat sensory neurons by a direct action. Neuroscience 145:686–698.

Verkest C, Piquet E, Diochot S, Dauvois M, Lanteri-Minet M, Lingueglia E, Baron A (2018) Effects of systemic inhibitors of acid-sensing ion channels 1 (ASIC1) against acute and chronic mechanical allodynia in a rodent model of migraine. British Journal of Pharmacology 175:4154–4166.

Wu B-M, Bargaineer J, Zhang L, Yang T, Xiong Z-G, Leng T-D (2020) Upregulation of acid sensing ion channel 1a (ASIC1a) by hydrogen peroxide through the JNK pathway. Acta Pharmacologica Sinica.

Xue Y, Ren J, Gao X, Jin C, Wen L, Yao X (2008) GPS 2.0, a tool to predict kinase-specific phosphorylation sites in hierarchy. Molecular & cellular proteomics: MCP 7:1598–1608.

Yan J, Wei X, Bischoff C, Edelmayer RM, Dussor G (2013) pH-evoked dural afferent signaling is mediated by ASIC3 and is sensitized by mast cell mediators. Headache 53:1250–1261.

Zeke A, Misheva M, Reményi A, Bogoyevitch MA (2016) JNK Signaling: Regulation and Functions Based on Complex Protein-Protein Partnerships. Microbiology and molecular biology reviews: MMBR 80:793–835.

Zelenka M, Schäfers M, Sommer C (2005) Intraneural injection of interleukin-1beta and tumor necrosis factor-alpha into rat sciatic nerve at physiological doses induces signs of neuropathic pain. Pain 116:257–263.

Zhou R, Wu X, Wang Z, Ge J, Chen F (2015) Interleukin-6 enhances acid-induced apoptosis via upregulating acid-sensing ion channel 1a expression and function in rat articular chondrocytes. International Immunopharmacology 29:748–760.

