## Supplementary Figures 1 and 2 for "C-Jun N-terminal kinase post-translational regulation of pain-related Acid-Sensing Ion Channels 1b and 3"

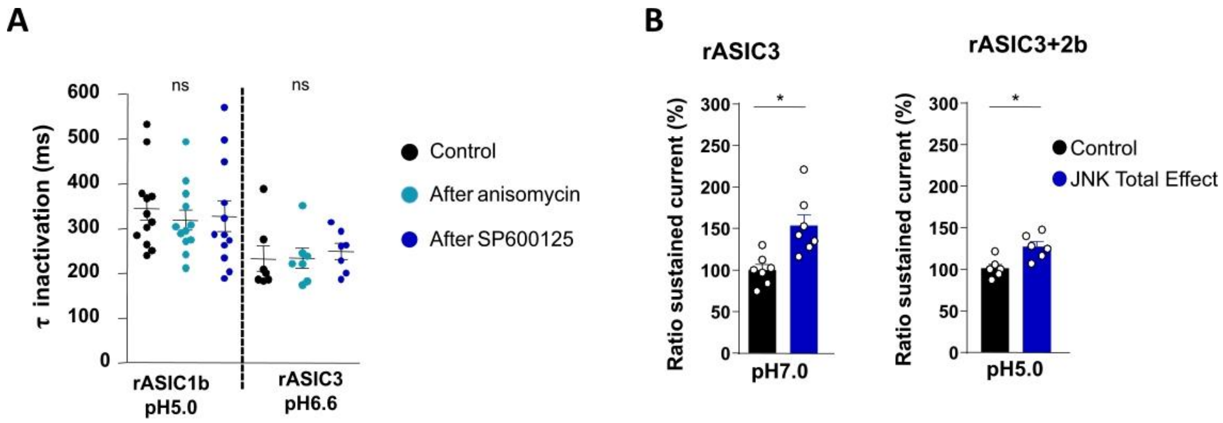


**<u>Supplementary Figure 1:</u> Effect of the JNK-regulation on inactivation time constants of rASIC1b and rASIC3 peak currents, and on the rASIC3 sustained currents**

**A,** Scatter plots of inactivation time constants (in ms) of rASIC1b peak current activated at pH5.0 and of rASIC3 peak current activated at pH6.6 from a conditioning pH of 7.4, before and after extracellular perfusion with anisomycin (50µM) and the JNK inhibitor SP600125 (50µM). One-way ANOVA ($F(2, 22)=0.43$, ns), n=7-11 cells. **B,** JNK-dependent potentiating effect on rASIC3 sustained window current activated at pH7.0 (left) and on rASIC3+2b pH5.0-evoked sustained current (right). Conditioning pH7.4, Wilcoxon paired test versus appropriate control, *, $p<0.05$, n=6-8. Total effect calculated and plotted as in Fig. 1B.

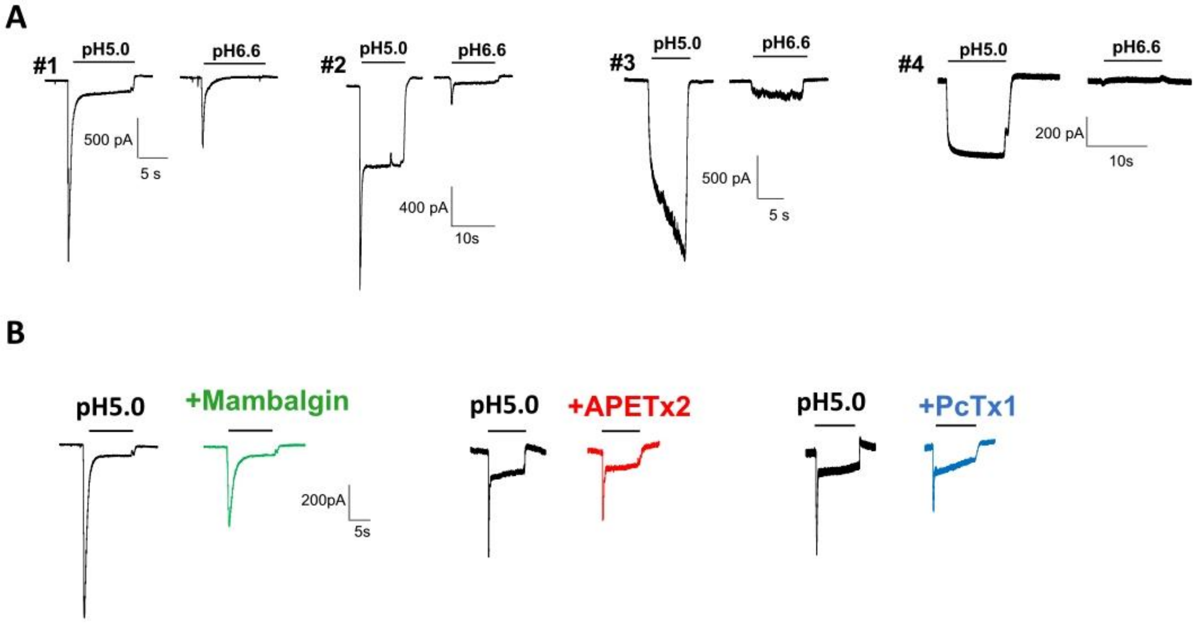


**<u>Supplementary Figure 2:</u> Characterization of ASIC-like currents in mouse DRG neurons**

**A**. Representative proton-activated inward currents recorded at -80mV after activation by pH5.0 and 6.6 from conditioning pH7.4 in cultured mouse DRG neurons from 4 different cells showing transient ASIC-like currents associated or not with a sustained current of variable amplitude (#1 and #2), or in neurons with only sustained currents without any peak (#3 and #4). **B**. Representative recordings from 3 different neurons showing at least 20% of inhibition of the ASIC-like currents by mambalgin-1 (1µM), APETx2 (3 µM) and PcTx1 (20 nM) applied in the conditioning pH7.4 solution.
